# A quantitative framework for bacterial competition during starvation

**DOI:** 10.64898/2026.05.19.726047

**Authors:** Zara Gough, Mara Dauber, Hamid Seyed-Allaei, Elena Biselli, Sophie Brameyer, Severin J. Schink, Ulrich Gerland

## Abstract

Bacterial communities often spend long periods under starvation, where survival depends not only on their intrinsic ability to withstand stress but also on nutrients released by dying neighbors. This creates a distinct form of competition: cells compete for recycled necromass, and the outcome should depend on physiological traits that determine nutrient uptake and maintenance demand. Here, we develop a quantitative framework for this competition using *Escherichia coli* populations whose starvation physiology is tuned by prior growth history. Fast-grown populations have higher maintenance demands and die slightly faster in monoculture, whereas slow-grown populations are better adapted to starvation. In co-culture, these physiological differences are strongly amplified in a frequency-dependent manner: less-adapted populations die several-fold faster than in monoculture, whereas well-adapted populations can reduce their death rate below that of stationary-phase adapted monocultures. We explain these dynamics with a shared-energy-pool model in which death releases recyclable nutrients, surviving cells consume them for maintenance, and intracellular energy sets death rate. Using independently measured parameters, the model makes parameter-free predictions for competitive survival. The predicted instantaneous death rates collapse onto a universal function of the population ratio over four orders of magnitude. Our results establish necromass recycling as a quantitative basis for bacterial competition during starvation and lay the foundation for modeling communities during starvation.

## Introduction

In nature, bacteria are selected not only for how fast they grow, but for how long they survive. Because bacteria proliferate rapidly when nutrients are available, phases of exponential growth tend to be brief; populations quickly exhaust accessible resources and spend extended periods under starvation. This phase, however, is not completely nutrient free. Once cells start dying of starvation, the remaining viable cells can use necromass released by dying neighbors as nutrients [43, 5, 39, 40]. If multiple species are starving as a community, this means that biomass from one population can be used by other populations, and as a consequence, survival—the central measure of fitness during starvation— transforms from an intrinsic property of individual cells into a frequency- and community-dependent variable. This implies that planktonic bacteria in nutrient-poor conditions interact as a community – something classically associated with nutrient-rich environments and high-density cultures [29, 8].

Arguably, starving bacterial communities are prevalent and important in nature. A canonical example is the pelagic ocean, where nutrient concentrations are generally very low, yet a large diversity of bacteria persist and compete for limiting resources [18]. These conditions favor specialized oligotrophic bacteria adapted to low nutrient availability such as SAR11 [17]. Host-associated environments are often considered nutrient rich, but bacteria within them nevertheless experience frequent transitions between growth and starvation due to strong spatial and temporal nutrient heterogeneity. Such fluctuations occur across diverse contexts, including infections [12], biofilms [44, 24], and the gut [33, 6, 10, 32].

Despite this broad relevance of starving communities, the quantitative toolkit of microbial ecology has largely focused on populations that are actively growing. Community dynamics during growth have been described using phenomenological approaches such as evolutionary game theory [31, 26, 47, 13, 30], Lotka–Volterra models [9, 28, 42], or pairwise survival rules [14]. More recently, consumer–resource models have provided a mechanistic framework for explicitly incorporating nutrient competition and cross-feeding [19, 23, 1]. A comparable mechanistic framework for competition during starvation is missing.

Developing such a framework is challenging for three reasons. First, necromass released by perished cells can be recycled by surviving community members [43], coupling each cell’s fate to the death of others. Even in monoculture, this coupling shapes starvation kinetics, producing approximately exponential viability decay in *Escherichia coli* [39] and longer-tailed decay in many other bacteria [40]. Second, key physiological parameters that determine starvation survival—such as maintenance demand and the ability to recycle necromass— not only vary across bacterial species [36, 46, 25], but even several-fold among isogenic cells depending on prior growth history [4, 38, 37]. Thus, it matters not only who competes, but also how competitors were physiologically primed before starvation. Third, bacteria can engage in active interference competition, including antimicrobial toxins [22, 21], phages [11, 27], and contact-dependent secretion systems [35, 45], which can further reshape survival by directly killing competitors and generating additional recyclable biomass.

This raises a simple yet unresolved question: when multiple starving populations share nutrients released from dead cells, how do the populations benefits from this necromass? In growing communities, competition is often framed in terms of nutrient uptake or growth yield. Under starvation, however, the relevant traits are different: cells must consume recycled nutrients for maintenance, and death itself supplies the common resource. We therefore asked whether competitive survival during starvation can be predicted from a small set of physiological parameters: death rate, maintenance demand, and uptake rates. Put differently, we ask whether the benefit of necromass released by dead cells can be predicted from measurable physiological traits of the surviving competitors.

We take a deliberately minimal approach to isolate this mode of competition. We use the well-defined dependence of death rate on prior growth rate in *E. coli* [4] and compete fast-grown against slow-grown populations that are genetically identical apart from a neutral antibiotic resistance marker. Prior growth history tunes starvation physiology: fast-grown cells have higher maintenance demand and die faster, whereas slow-grown cells are better adapted to starvation [4]. This design removes genetic and metabolic confounders and lets us ask whether recycling interactions alone can generate predictable selection under starvation.

We find that co-culture survival deviates strongly from the null expectation based on isolated monocultures. Death rates shift several-fold in either direction depending on population ratio, and these shifts are quantitatively predicted by a shared-energy-pool model in which dying cells release recyclable nutrients and surviving cells consume them for maintenance. The central prediction is that starvation competition is governed by two physiological levers: uptake determines how much of the common pool a population accesses, while maintenance demand determines how much energy is required to reduce death rate. Strikingly, selection under starvation competition is frequency dependent and favors strains already better adapted to starvation, thereby amplifying competitive differences and destabilizing community composition.

## Results

### Co-cultured populations show ratio-dependent shifts in death rates

To probe competition under starvation in a minimal setting, we designed co-culture experiments using *E. coli* populations that were genetically identical except for neutral antibiotic resistance markers (Table S1). The populations differed only in their physiological state, set by prior growth history. Specifically, cells were pre-grown in mannose (slow), glycerol (moderate), or glucose (fast), and then transferred into carbon-free minimal medium.

Previous work has shown that the decrease of viability in minimal medium is approximately exponential [39, 34], and that faster prior growth increases maintenance demand due to proteome-partitioning affecting the cell’s ability to decrease its permeability [4, 37, 38]. This higher maintenance rate leads to faster death during starvation [4]. Consistent with these expectations, we observed an approximately exponential decline in viability over 7 days, measured by daily plating on LB agar. The death rate depended on prior growth rate, with glucose-grown cultures dying about twice as fast as mannose-grown cultures (Fig. 1B–C).

**Figure 1.**
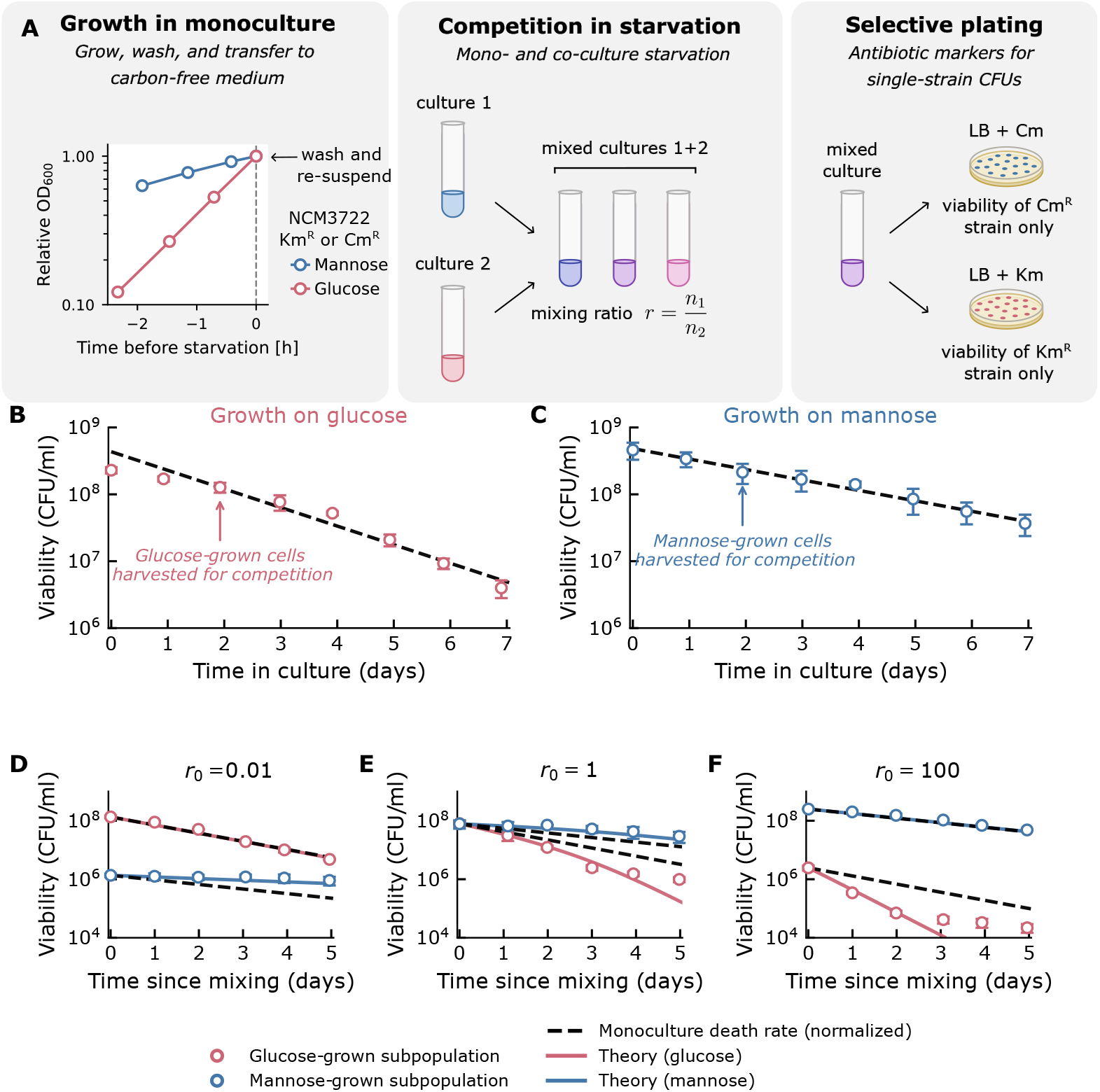
(A) Schematic of the experiment. (B,C) Monoculture viability of glucose-grown and mannose-grown cultures; circles denote experiments (*±*1 s.d.), black dashed lines are exponential fits; arrows mark the sampling time (*∼*2 days) used to seed the competition assays below. (D–F) Glucose–mannose competitions at initial mixing ratios *r*_0_ = 0.01, 1, and 100 (mannose:glucose). Circles show experiments (*±*1 s.d.); solid colored lines are the parameter–free theory for each mixture; black dashed lines reuse the same monoculture fit from panels B–C, rescaled to the initial viability of each competition. See Fig. S1 for more competition experiments.

When populations with different prior growth histories were mixed, the resulting survival kinetics deviated strongly from monoculture death rates (Fig. 1D–F). For example, when mannose- and glucose-grown populations were mixed such that the glucose-grown subpopulation was in the majority (and mannose-grown in the minority), the death rate of the mannose-grown subpopulation decreased by about a factor of three relative to its monoculture value (Fig. 1D), leading to a total six-fold slower death rate than the glucose-grown culture. Conversely, when the glucose-grown subpopulation was in the minority, its viability declined approximately threefold faster compared to its monoculture reference (black dashed line), while the mannose-grown subpopulation decayed at a rate similar to the monoculture (Fig. 1F). At intermediate mixing (near equal abundances), both death rates shifted to intermediate values (Fig. 1E).

These trends were robust across replicate experiments, yet sensitive to both the initial mixing ratio and the pre-culture condition (Fig. S1). The more a strain was in the minority, the more strongly its death rate shifted away from its monoculture value; in contrast, when a strain was in the majority, its dynamics approached monoculture behavior (black dashed lines in Fig. 1 and Fig. S1). Together, these findings reveal that co-cultures do not simply superimpose the survival properties of their constituent populations, but instead amplify physiological differences in a frequency-dependent manner.

### A shared energy-pool model for competition in starvation

In monoculture, starvation death is quantitatively set by a balance between nutrient release from dying cells and nutrient consumption by surviving cells [39, 4, 38]. As cells die, they release biomass that can be recycled by the remaining population and used for maintenance. Populations with higher maintenance requirements consume recycled nutrients faster, enter a lower-energy internal state, and therefore exhibit higher death rates. The higher death rates lead to more nutrients becoming available for the surviving cells, which they can use to meet their maintenance requirements. This feedback self-adjusts until nutrient release through death balances nutrient consumption by survivors, causing the population death rate to become quantitatively determined by the ratio between maintenance demand and recycling yield.

Guided by this logic, we built a framework that explicitly links recycling from dying cells to the maintenance and survival of their neighbors via a shared extracellular energy pool, see Fig. 2.

**Figure 2:**
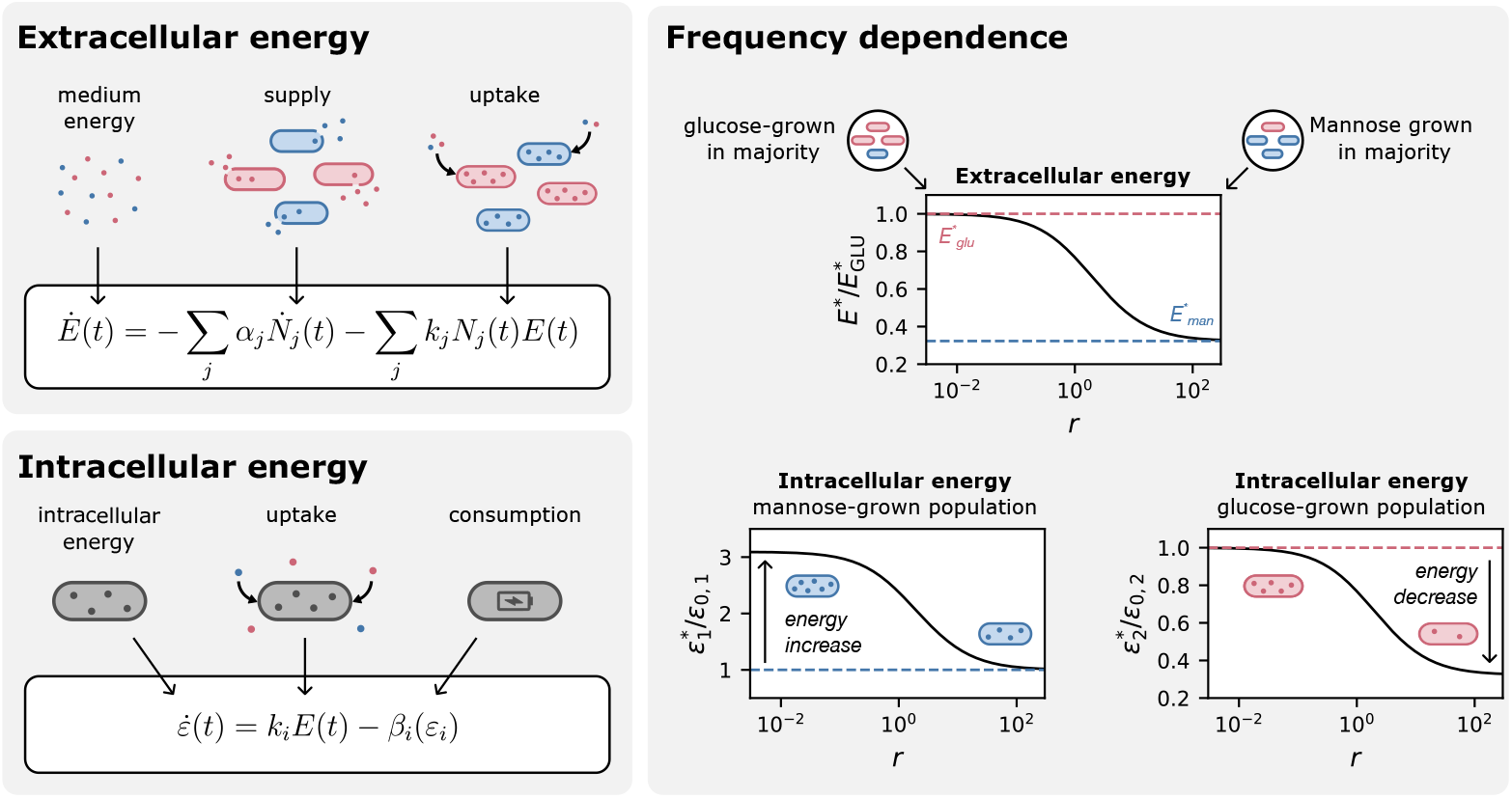
Extracellular energy. The nutrient pool in the supernatant is supplied with necromass released by dying bacteria and consumed by all bacteria in the community. **Intracellular energy** The energy pool inside bacteria of each subpopulation is supplied by nutrient uptake and consumed by maintenance processes. **Frequency dependence** The extracellular energy pool depends on the ratios of subpopulations 1 and 2. The internal energies respond to the extracellular energy pool and therefore deviate several-fold from the monoculture internal energy states in the minority states (low r for subpopulation 1 and high r for subpopulation 2), leading to changes in death rate.

We consider any number of competitors indexed by *j* = 1, …, *J*. We approach this by first determining how much energy is available to the community (the extracellular pool), and then determining how much of that energy an individual cell uses (its consumption flux).

#### Extracellular energy balance

The shared extracellular energy pool, *E*(*t*), quantified as the total energy in the culture volume, increases by release of nutrients from dying cells and decreases as viable cells take up these nutrients. We assume that uptake is pool-limited and unsaturated, i.e. proportional to *E*. The flux balance is expressed by

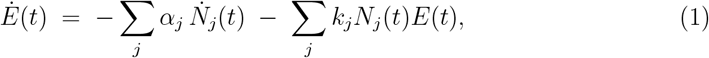

where *α*_*j*_ is the recycling yield (i.e., the amount of energy that can be obtained from a dead cell), *k*_*j*_ the uptake rate, and *N*_*j*_(*t*) the number of viable cells at time *t* (with 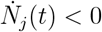 during starvation).

#### Intracellular energy balance

For a viable cell of strain *i*, intracellular energy *ε*_*i*_(*t*), quantified by the total energy available inside a cell, changes by uptake and consumption,

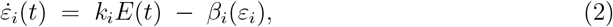

with *β*_*i*_(*ε*_*i*_) the consumption rate.

Because nutrient fluxes usually equilibrate rapidly compared to the slow decay of viability (days), we assume a quasi–steady state for both extracellular and intracellular pools.

Setting 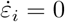 gives

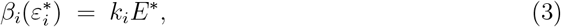

and *Ė*= 0 in Eq. (1) yields

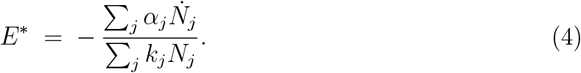

Combining these relations determines each strain’s steady consumption flux as its share of the total release:

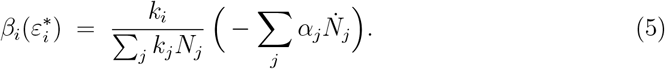

Equation (5) implies that each cell accesses a fraction of the recycled nutrients proportional to its uptake capacity *k*_*i*_ relative to the community total ∑_*j*_ *k*_*j*_*N*_*j*_. This partitioning result constitutes the basis for linking consumption to intracellular energy and, in turn, to death rates in mixed cultures.

### Intracellular energy links consumption and death

To close the model, we connect intracellular energy to the physiological processes of consumption and survival in the low-energy, non-growing regime characteristic of starvation, exploiting established constraints and experimental evidence.

Consumption rate *β*_*i*_(*ε*_*i*_) must vanish when no energy is available and should increase monotonically with intracellular energy. The simplest plausible functional form is a linear expansion about *ε* = 0:

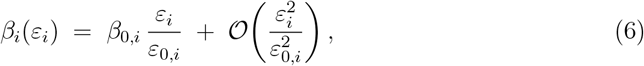

where *β*_0,*i*_ is a reference consumption rate (e.g. in monoculture) and *ε*_0,*i*_ the corresponding intracellular energy scale in that state for strain *i*. Higher-order terms may appear (e.g., if energy becomes sufficient to sustain growth), but the linear contribution is the leading-order term at low energy.

For the death rate *γ*_*i*_(*ε*_*i*_), recent experiments show two key features. First, if extracellular nutrients are constantly washed away death rate accelerates with an exponentially increasing rate, consistent with a Gompertz law [48], i.e., a divergence as *ε* → 0 in our picture. Second, nutrient supplementation can arrest death on experimental timescales [39], demonstrating that sufficiently large *ε* stabilizes survival. The simplest function capturing both behaviors is hyperbolic:

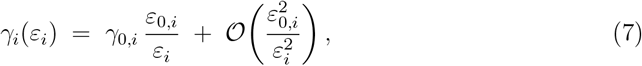

where *γ*_0,*i*_ is the monoculture reference death rate.

Together, Eqs. (6) and (7) lead to a reciprocal relationship between consumption and death in quasi-steady state. As intracellular energy decreases, consumption falls while death accelerates; conversely, when consumption sustains intracellular energy, death slows.

At quasi–steady state, this reciprocity reads

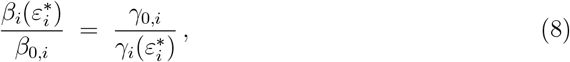

so that a higher consumption rate relative to its monoculture reference corresponds directly to a lower death rate, and vice versa.

### Model prediction of death rates in co-culture

We now derive the death rates of competing strains from the coupled energy-pool relations. For a general community with *J* competitors, the quasi–steady balances yield a closed system for the *γ*_*j*_; the full derivation is provided in the Supporting Information (SI, *Derivation of the Model*).

For the two–strain case that we studied experimentally, it is convenient to express the solutions in terms of the viability ratio *r* ≡ *N*_1_*/N*_2_. Eliminating *E*^*\**^ and 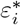 via the quasi–steady relations and using the consumption–death reciprocity (Eq. (8)) gives:

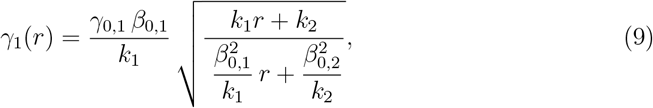

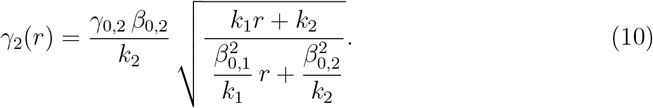

These equations capture the central logic of competition during starvation survival. Bacteria can reduce their death rate in two fundamentally distinct ways: by increasing nutrient uptake, thereby securing a larger share of the common energy pool to sustain maintenance, or by lowering their maintenance demand, allowing them to survive longer on the same energetic share. Importantly, lowering maintenance demand benefits a strain even without directly altering nutrient release or uptake kinetics: because all cells draw from the same shared energy pool, a strain that requires less maintenance can indirectly profit from the death of its competitors and thereby reduce its own death rate. Next, we will quantify the parameters used in Eqs. (9)–(10) to check what determines the competition outcomes in our *Escherichia coli* experiments.

### Physiological parameterization and model validation

The death rates of Eqs. (9)–(10) depend on the monoculture consumption rates *β*_0,*i*_, and the uptake rates *k*_*i*_ and the recycling yield *α*_*i*_ (via the monoculture death rates *γ*_0,*i*_ = *β*_0,*i*_*/α*_*i*_). These parameters are measurable and allow us to obtain parameter–free predictions for competitive survival kinetics under starvation.

The monoculture death rate *γ*_0_ is measured in parallel for each pre-growth condition (Fig. 1B–C). The consumption (maintenance) rate *β*_0_ is taken from Biselli *et al*., who report an approximately exponential increase with prior growth rate *µ* [4, 3],

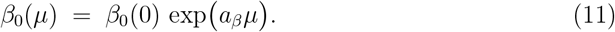

with *a*_*β*_ ≈ 2.1h.

The remaining component is the uptake capacity *k*. Because uptake is pool-limited and unsaturated in our framework, we assume it scales with transport area. Using cell size characterizations from Biselli *et al*. [4], we compute the membrane surface area and parameterize it also as an exponential in *µ*:

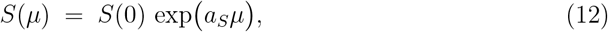

with *a*_*S*_ ≈ 0.25h and set *k*(*µ*) *∝ S*(*µ*) (only ratios of *k* enter Eqs. (9)–(10)).

Inserting *γ*_0_ (measured), *β*_0_(*µ*) (from Biselli et al), and *k*(*µ*) (surface area calculated using geometry data from Biselli et al) into Eqs. (9)–(10) yields parameter-free predictions for competitive death rates. These appear as the solid colored lines in Fig. 1D–F and Fig. S1, capturing fast-, moderate-, and slow-growing physiological states across the full range of initial mixing ratios. The only consistent deviation is a flattening of the viability curves for fast-dying subpopulations (e.g., Fig. 1F), which we analyze later.

A test of the theory is to view the instantaneous death rate *γ*_*i*_(*t*) as a function of the contemporaneous viability ratio *r*(*t*) = *N*_1_(*t*)*/N*_2_(*t*). Despite the added noise from estimating *γ*_*i*_(*t*) over short 1–2 day windows (Methods), curves from all initial mixtures *r*_0_ collapse onto a single master curve *γ*_*i*_(*r*) with no adjustable parameters (Fig. 3A–B). This collapse is precisely what Eqs. (9)–(10) predict: for fixed physiological parameters, each death rate depends only on the single scaling variable *r*. The agreement holds over four orders of magnitude in *r* (10^*−*2^–10^2^), smoothly connecting the minority and majority limits (dashed lines), and is reproduced for the other substrate pairs (SI Fig. S2).

**Figure 3:**
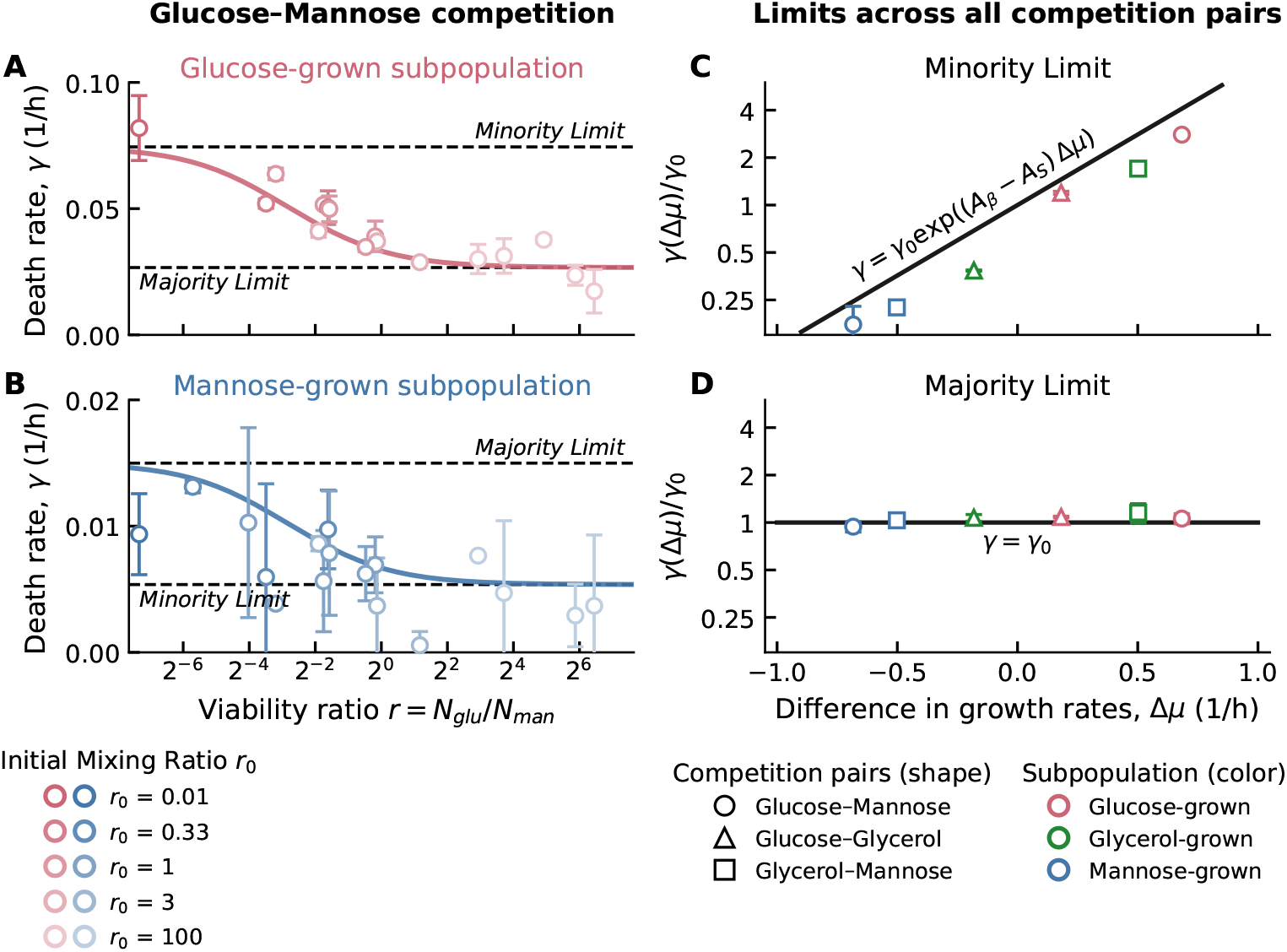
Death rates in two–strain competitions and theory. **A–B (left):** Measured instantaneous death rates *γ* (1/h) versus the ratio *r* = *N*_man_*/N*_glu_ for glucose–mannose competitions, shown separately for the glucose-grown (A) and mannose-grown (B) subpopulations. Points are mean *±* s.d.; different shades denote different *r*_0_ values. The solid curve is the model prediction from Equations 9 and 10. Dashed horizontal lines mark the theory’s minority and majority limits. **C–D (right):** Normalized death rate *γ*(Δ*µ*)*/γ*_0_ versus the difference in growth rates Δ*µ* (1/h), using experiments started near the minority limit (*r*_0_ *<* 0.01, panel C) and majority limit (*r*_0_ *>* 100, panel D); *γ* is estimated over the full starvation period. Marker shape indicates the competition pair (glucose–mannose, glucose–glycerol, glycerol–mannose) and color indicates the subpopulation (glucose-, glycerol-, or mannose-grown). The black line in C is the minority-limit prediction (Eq. (14)); the black line in D is the majority-limit prediction *γ/γ*_0_ = 1 (by definition of *γ*_0_, Eq. (13)). Datasets for glucose–glycerol and glycerol–mannose are described in the Supporting Information.

### Minority and majority limits reveal an energy-pool mechanism

We analyze the limiting behaviors of the two–strain solutions to identify what sets death rates when one strain is rare or dominant. From Eqs. (9)–(10) it follows that each majority behaves like its monoculture,

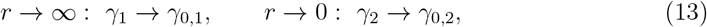

whereas in the minority limit *N*_min_ ≪ *N*_maj_ the rare strain’s death rate reduces to (SI Eq. S25)

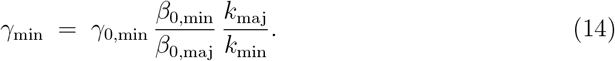

These limits correspond to the dashed “minority/majority” lines in Fig. 3A–B and extend to the other substrate pairs (SI Fig. S2). When compared to death rates measured across multiple time points, we see that these predictions match quantitatively in Fig. 3C (minority limit) and Fig. 3D (majority limit).

Using the empirical dependencies 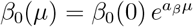 and 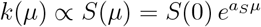 yields

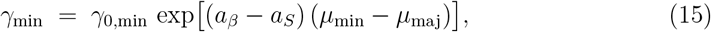

he straight line in Fig. 3C. The sign of Δ*µ* = *µ*_min_ *− µ*_maj_ determines the direction: Δ*µ >* 0 (fast-grown minority, e.g., glucose) increases the death rate 3-fold, whereas Δ*µ <* 0 (slow-grown minority, e.g., mannose) decreases it 3-fold. These two cases are two sides of the same exponential law.

Because *a*_*β*_*/a*_*S*_ ≈ 8, most of the predicted change in death rate of *Escherichia coli* is due to changes of the maintenance rate, rather than uptake rate. This shows that maintenance rate can be a major driver in competition for necromass.

Although the model quantitatively captures starvation dynamics across competition experiments, systematic deviations appear for fast-dying populations: after several days, the decline in viability slows and the apparent death rate flattens. This behavior corresponds to a biphasic decay, which is also observed in monocultures once viability has fallen to approximately 1% of the initial population (Fig. S3).

Such biphasic survival curves suggest the presence of a long-lived subpopulation with a lower effective death rate. This subpopulation may arise from preexisting phenotypic heterogeneity, such as a non-growing persister fraction [2], or from a starvation-induced physiological transition during prolonged nutrient deprivation. These long-lived survivors mark the transition into long-term stationary phase (LTSP) [15, 43].

Long-lived survivors isolated at the end of the experiment showed similar biphasic dynamics in repeat experiments (Fig. S3), arguing against a mutational origin. Instead, the data indicate that even genetically uniform monocultures can spontaneously generate coexisting subpopulations with distinct survival capacities. In this sense, the observed late-time deviation does not contradict the model, but rather mirrors its central principle: survival dynamics emerge from interacting subpopulations with different effective death rates.

## Discussion

Starvation survival is a central fitness trait of bacteria, alongside growth and reproduction. Here we show that survival in co-culture is governed by a simple, quantitative principle: a *necromass recycling feedback* that links nutrient release from dying cells, uptake by survivors, and maintenance demand. This feedback makes death rates of competing sub-populations interdependent and predictable from a small set of physiological parameters that scale with growth history.

A simple way to read the model is to start from the shared pool *E*^*\**^: it is the common “reservoir” of recyclable nutrients in the supernatant (Eq. (4)). Because uptake is pool-limited, each cell’s consumption flux is proportional to that pool, 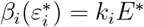 (Eq. (3)). Intracellular energy 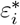 rises when this flux is high and falls when it is low; the death rate 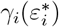 decreases with increasing 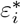 (Eq. (7)). Put together, the external pool *E*^*\**^ effectively sets the death rate: big pool ⇒ higher *ε* ⇒ lower *γ*; small pool ⇒ lower *ε* ⇒ higher *γ*.

Competition is then defined by the factors that fix *E*^*\**^. The energy pool is continually filled by the release from dying cells and drained by the uptake of survivors (Eq. (4)). If the majority dies slowly, little energy is released, so *E*^*\**^ remains low; if the majority dies fast, a lot of energy is released, and *E*^*\**^ rises. A minority strain therefore experiences a pool that is typically different from its own monoculture: a fast-grown (high-maintenance) minority faces a lower *E*^*\**^ than it would in monoculture and thus dies faster; a slow-grown (low-maintenance) minority receives a higher *E*^*\**^ than it would alone and thus dies more slowly.

The minority-limit formula (Eq. (14)) makes the two levers explicit. The ratio *β*_0,min_*/β*_0,maj_ captures the maintenance demand (the flux required by the minority to sustain its *ε*), and *k*_maj_*/k*_min_ captures the relative uptake capacity of majority versus minority populations (the inverse of the share of the energy pool consumed by the minority). Substituting the empirical growth-history scalings 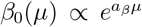 and 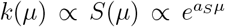 into Eq. (14) gives the exponential law (Eq. (15)). The sign of Δ*µ* = *µ*_min_ *− µ*_maj_ then immediately explains both outcomes we observe experimentally: a glucose minority (Δ*µ >* 0) dies about threefold faster than in monoculture, whereas a mannose minority (Δ*µ <* 0) dies about threefold slower (Fig. 3C–D).

### Motivation for a minimal competition scenario

Rather than comparing different species, we designed a two-player competition scenario between nearly isogenic *E. coli* strains that differ only in their physiological state set by prior growth (and a neutral antibiotic marker; neutrality confirmed in Table S1). This design removes genetic and regulatory confounding factors and reduces competitive survival to three experimentally grounded parameters in monoculture: the death rate *γ*_0_ (measured here, Fig. 1B–C), the consumption rate *β*_0_(*µ*) (approximately exponential in *µ*; Eq. (11)), and the uptake rate *k*(*µ*) inferred from cell’s surface area *S*(*µ*) (also exponential; Eq. (12), with *k* ∝ *S*). Under starvation, where uptake is pool-limited and nutrients arise solely from recycling, inserting these growth-history dependencies into Eqs. (9)–(10) yields parameter-free predictions. Empirically, instantaneous death rates from mixtures with different initial ratios collapse onto a single master curve *γ*_*i*_(*r*) over four orders of magnitude in *r* (Fig. 3A–B), and the minority/majority limits of the model align with experimental data (Fig. 3C–D, see also SI Fig. S2). This experimental design with minimal genetic differences between competitor strains shows that phenotypic (physiological) differences alone are sufficient to produce and to predict selection under starvation.

### Broader implications

Natural communities consisting of several distinct bacterial species differ not only in maintenance demand *β*_0_ and uptake capacity *k*, but also in which recycled molecules they can use, how they transport them (distinct transporters and regulation), and whether they directly antagonize competitors (bacteriocins, antibiotics, toxin–antitoxin systems, phages, etc.). Our minimal, isogenic design therefore captures a baseline mode of starvation competition that should be accounted for before invoking niche preferences or warfare. Ignoring this baseline risks conflating intrinsic maintenance/uptake asymmetries with substrate specificity or interference effects. The framework is readily extensible: one can introduce substrate-specific recycling and uptake (vectors ***α*** and ***k***), weak uptake saturation, or additional mortality/production terms to model antagonism. In many starved settings, the recycling term may provide the baseline contribution, with nutrient specificity and interference occurring as additional effects.

Overall, by linking prior growth history to maintenance and uptake, and by tying both to a shared resource pool, we provide a mechanistic and predictive basis for selection during starvation. This vantage point should be useful for analyzing multispecies competitions, structured environments such as biofilms, and evolutionary trade-offs between rapid growth and long-term survival.

## Author Contributions

**Conceptualization:** ZG, EB, SJS, UG

**Methodology:** ZG, HSA, EB, SJS, UG

**Investigation:** ZG, MD; with assistance from HSA

**Resources (strain construction):** SB

**Formal Analysis / Modeling:** ZG, SJS, UG; with assistance from HSA

**Writing – Original Draft:** ZG, SJS, UG

**Writing – Review & Editing:** All authors

**Supervision:** SJS, UG

*SJS and UG are co-corresponding authors and contributed equally as co-senior authors*.

## Declaration of Interests

The authors declare no competing interests.

## Methods

### Strains

The bacterial strains used in this study are derived from wild-type *E. coli* K-12 strain NCM3722 [41].

The chloramphenicol-resistant strain, NCM3722-*cat*, was made by double-homologous recombination using the suicide plasmid, pNPTS138-R6KT-*cat*. Two DNA fragments comprising 650 bp as the flanking region of the attachment site were amplified by PCR using *E. coli* NCM3722 genomic DNA as a template. The gene *cat* with its promoter was amplified by PCR using the plasmid pBAD33 as a template [20]. After purification, these three fragments were assembled via Gibson assembly [16] into EcoRV-digested pNPTS138-R6KT plasmid, resulting in the pNPTS138-R6KT-*cat* plasmid. This plasmid was introduced into *E. coli* NCM3722 by conjugative mating using *E. coli* WM3064 as a donor in LB containing 300 µM meso-diamino-pimelic acid (DAP). Single-crossover integration mutants were selected on LB plates containing kanamycin but lacking DAP. Single colonies were grown over a day without antibiotics and plated onto LB plates containing 10% (w/v) sucrose but lacking NaCl to select for plasmid excision. Kanamycin-sensitive but chloramphenicol-resistant colonies were checked for targeted insertion by colony PCR using primers bracketing the site of the insertion. Insertion of *cat* was verified by colony PCR and sequencing. The genes *neo* and *cat* each were inserted with their individual promoter at the attachment site (between *glmS* and *rpmE*) of *E. coli* NCM3722.

The kanamycin-resistant strain, NCM3722-*neo*, was constructed via *λ*-Red recombination, where the rpsL-*neo* fragment was used as a template following the protocol of the “Quick and Easy *E. coli* Gene Deletion Kit” (Gene Bridges, Heidelberg). Insertion of *neo* was verified by colony PCR and sequencing.

### Culture Media

The minimal culture medium for *E. coli* used throughout this study is based on N-C-culture medium [7], containing 4 g/L K2SO4, 54 g/L K2HPO4, 18.8 g/L KH2PO4, 0.1 g/L MgSO4.7H2O and 10 g/L NaCl. 20 mM NH4Cl was added as a nitrogen source. The carbon sources used were 0.01% mannose, 5 mM glycerol and 0.05% glucose. All chemicals were purchased from Carl Roth, Karlsruhe, Germany.

### Growth

Cells were taken from a -80^*°*^C LB/glycerol stock and streaked on an LB agar plate, then incubated for around 16 h at 37^*°*^C. A single colony was picked with an inoculating needle and grown on pre-warmed LB for 3-4 h at 37^*°*^C as a seed culture. The culture was diluted and re-inoculated in pre-warmed minimal medium with either mannose, glycerol, or glucose, then re-diluted after growing exponentially for several doublings to form the experimental culture. Experimental cultures underwent at least 3 doublings prior to measuring growth. Growth rate was determined by measuring optical density using a spectrophotometer (*λ*=600 nm).

All growth and starvation took place in a water bath shaker (WSB-30, Witeg, Wertheim, Germany) at 250 rpm with water bath preservative (Akasolv, Akadia, Mannheim, Germany). Small cultures of 3 mL, 5 mL and 10 mL were grown in 18 *×* 150 mm, 20 *×* 150 mm and 25 *×* 150 mm disposable glass test tubes (Fisher Scientific, Hampton, NH, USA) respectively with disposable, polypropylene Kim-Kap closures (Kimble Chase, Vineland, NJ, USA). Larger culture volumes of 50 mL and 100 mL were grown in 250 mL and 500 mL baffled Erlenmeyer flasks (Carl Roth, Karlsruhe, Germany) with Kim-Kap closures.

### Starvation

Sudden starvation was induced by centrifuging cultures during exponential growth for 5 min at 3000 rcf at 37^*°*^C. The supernatant was removed and replaced with fresh, pre-warmed minimal medium without a carbon source.

Viability was measured approximately every 24 h by plating serially diluted samples of starving culture on LB agar and counting colony-forming units (CFU) after incubation at 37^*°*^C for approximately 18 hours. Samples were diluted in minimal medium without carbon substrate and spread on LB agar using Rattler Plating Beads (Zymo Research, Irvine, CA, USA). LB agar was supplemented with 25 *µ*g/mL 2,3,5-triphenyltetrazolium chloride to stain colonies and increase contrast for automated counting (Scan 1200, Interscience, Saint-Nom-la-Breteche, France). Samples were diluted such that 100-300 colonies grew on each petri dish (92 x 16 mm, Sarstedt, Nümbrecht, Germany). Staining or automation of counting had no significant effect on viability measurements or accuracy, compared to unstained, manually counted samples (*<*1% systematic error).

### Competition

Cultures with two different antibiotic markers were grown on two different sugars as described in ‘Growth’, then starved for 48 hours prior to the initiation of competition experiments. The two starving cultures were combined in ratios of 1:99, 50:50 and 99:1, or 75:25, 50:50 and 25:75, where the ratio refers to the number of viable cells of each respective strain at the initiation of competition. The required volumes of each culture were based on the viability measurement at 24 h and the expected death rate of each sugar, with a final total volume of 3 mL for each mixed-strain culture. Viability of each strain was measured as described in ‘Starvation’ – each sample was plated on a set of 3 LB agar plates containing 50 *µ*g/mL kanamycin, and a set of 3 LB plates containing 25 *µ*g/mL chloramphenicol, in order to determine the viability of each individual strain.

## A Supplementary Figures

**Figure S1:**
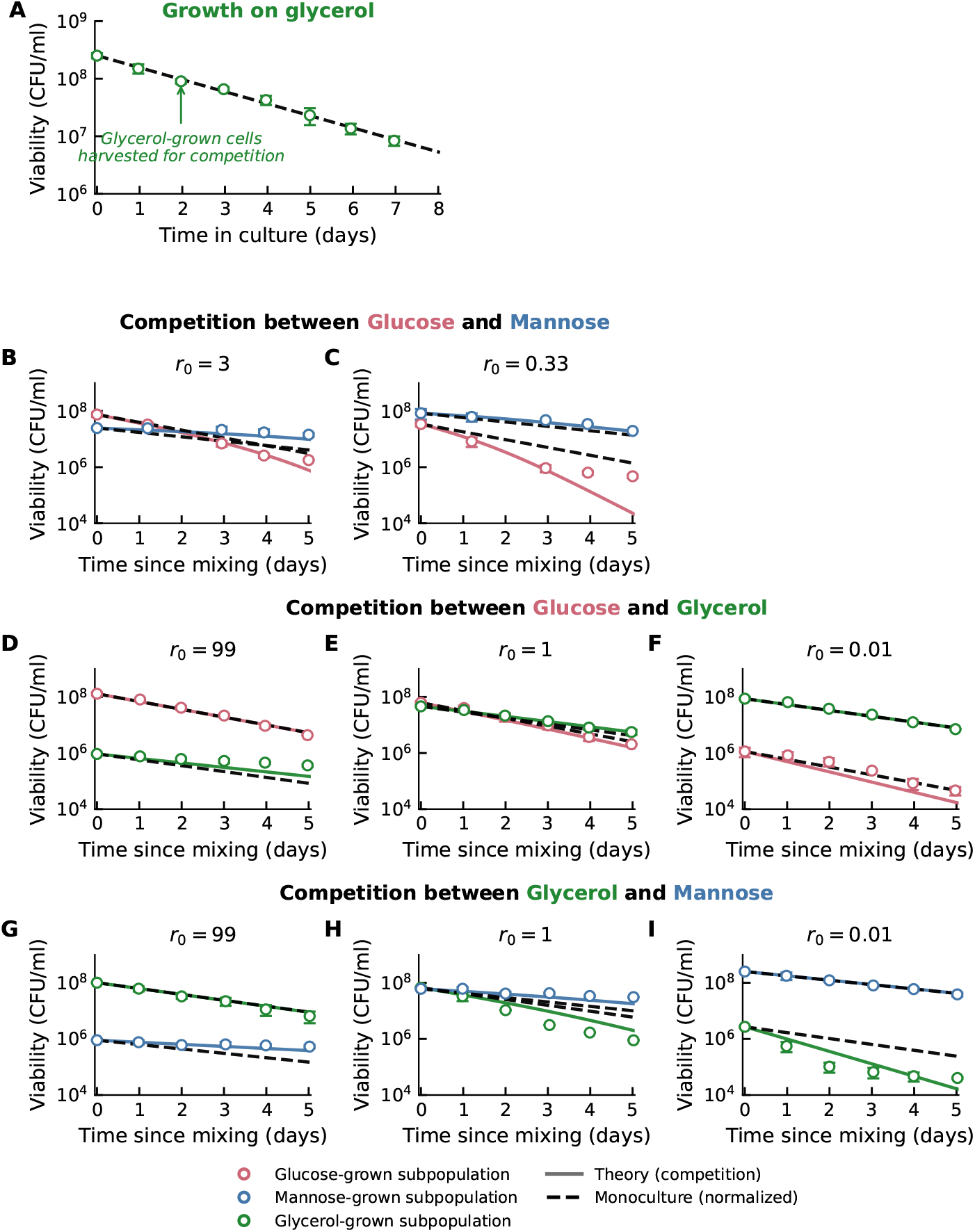
Supporting Figure S1. Additional monoculture and competition starvation datasets showing viability over time. **Row 1 (A):** Glycerol monoculture (glycerol-grown cells). Circles show experiments (*±*1 s.d.); the black dashed line is the corresponding monoculture theory. **Row 2 (B–C):** Glucose–mannose competitions (complementary to Fig. 1); initial mixing ratios *r*_0_ are indicated in each panel title. **Row 3 (D–F):** Glucose–glycerol competitions. **Row 4 (G–I):** Glycerol–mannose competitions. In all competition panels, circles denote experiments (*±*1 s.d.), solid colored lines are the parameter-free mixture theory, and black dashed lines reuse the same monoculture theory (for the matching subpopulation) rescaled to the initial viability of each competition (as in Fig. 1). Marker colors indicate subpopulation identity: glucose (red), glycerol (green), mannose (blue).

**Figure S2:**
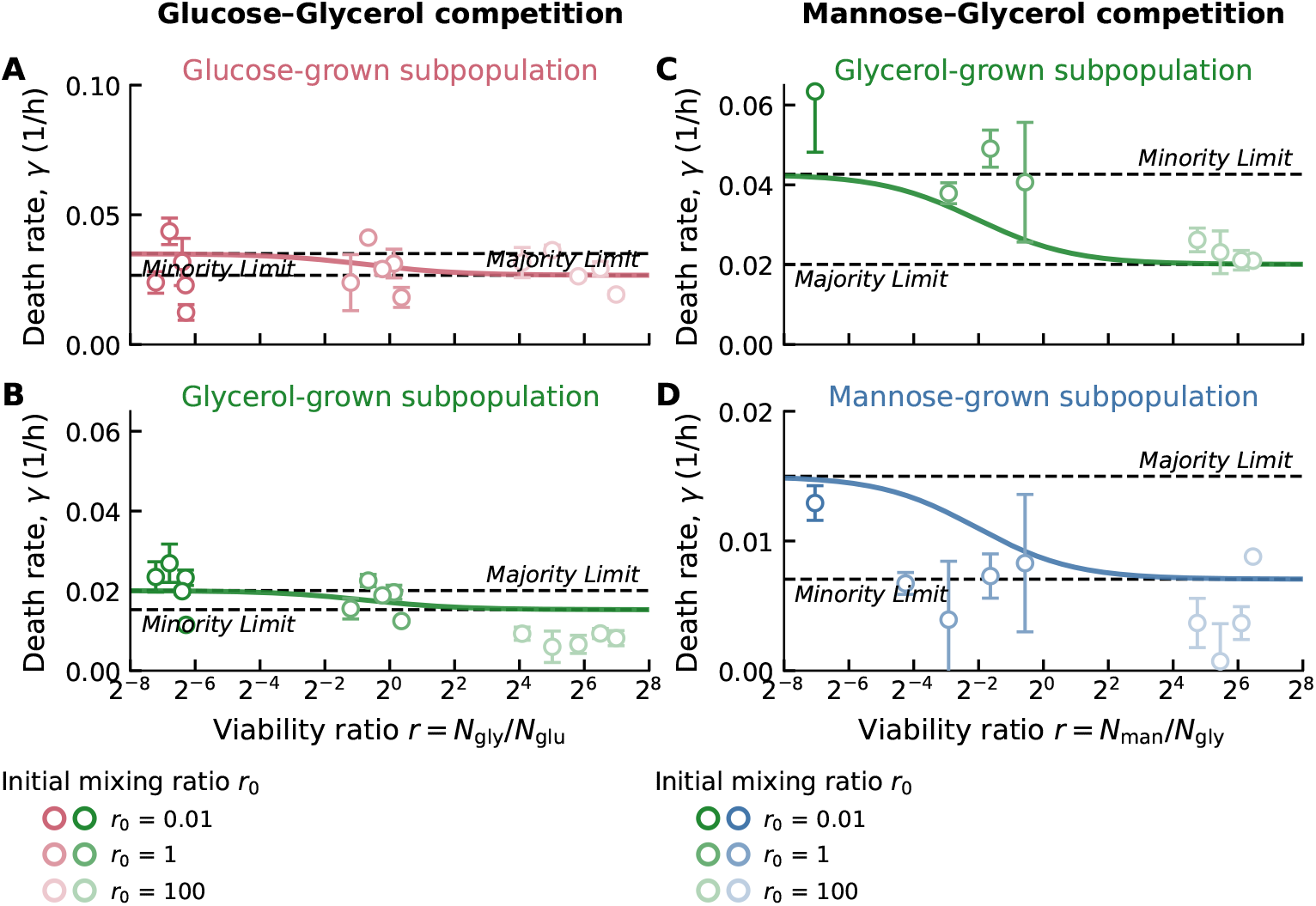
Death-rate amplification in *glucose–glycerol* (left column) and *mannose–glycerol* (right column) competitions, plotted in the same style as Fig. 3. Y-Axes have the same scale for each subpopulation across Fig.1 and this figure. Top panels show the subpopulation indicated in each title; bottom panels show the partner subpopulation from the same competitions. Symbols are experimental measurements at different initial mixing ratios *r*_0_; shading encodes *r*_0_, so the two subpopulations from the same initial mixture share the same shade. In each panel the abscissa is the viability ratio *r* = *N*_carbon1_*/N*_carbon2_ (see axis labels). Solid curves are the parameter-free predictions of the model from the main text (Eqs. (9) and (10)). Horizontal dashed lines denote the minority and majority limits implied by the same theory (Eqs. (14) and (13) respectively) together with the physiological scalings in Eqs. (11)–(12).

**Figure S3:**
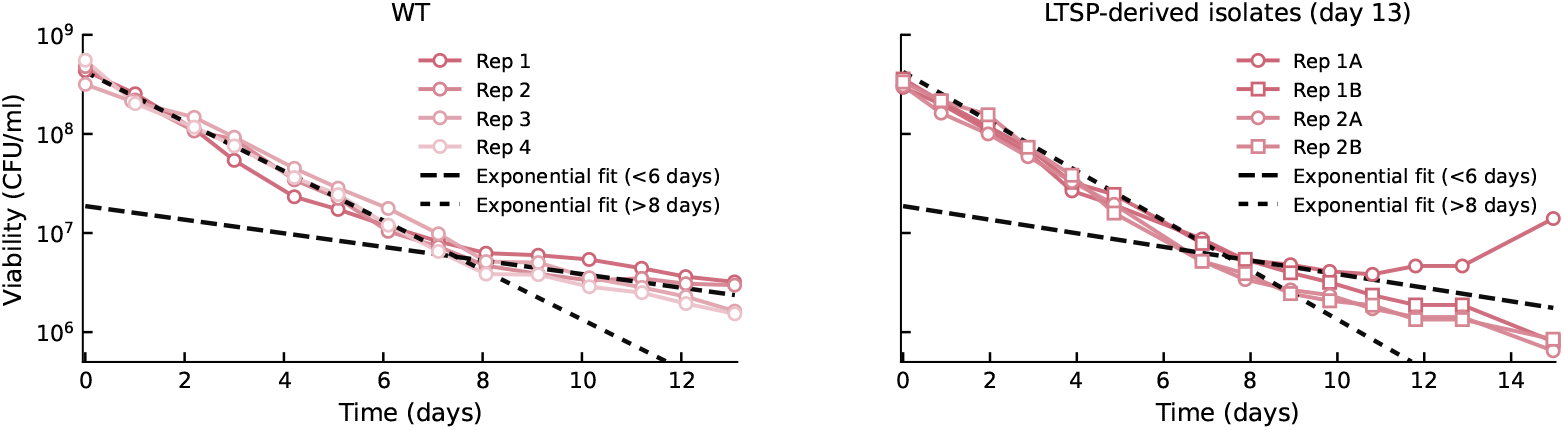
Biphasic decay and transition to long-term stationary phase. Left. Four replicates of glycerol-grown *E. coli* starving in monoculture (Rep 1–4) measured for 13 days. Black dashed curves indicate exponential fit for an early (first six days) and a late (after eight days) phase, showing the presence of a biphasic decay. This phase is typically followed by long-term stationary phase [43]. Right. Colonies picked at the end of the experiment Rep 1 and Rep 2 were regrown in minimal medium and starved identically as in the original experiment. Survival curves again showed a biphasic decay that was similar to the original wild-type.

## Derivation of the Model

Here we derive the mathematical model that links nutrient recycling, intracellular energy, and death rates during starvation. The logic proceeds in several steps. We begin with the extracellular energy balance, which describes how nutrients are released by dying cells and taken up by surviving cells. We then derive the intracellular energy balance and, under a quasi–steady assumption, express per–cell consumption in terms of community release and uptake. We next connect consumption to intracellular energy and assume that the death rate depends on intracellular energy. Substituting the quasi–steady relations closes the loop, yielding coupled equations for the death rates of all strains. Finally, we analyze limiting cases to verify that the model recovers monoculture behavior and gives consistent predictions for mixed cultures.

### Core assumptions

(i) Uptake is pool-limited and unsaturated: uptake flux is proportional to the extracellular energy pool *E*. (ii) External and internal energy pools typically equilibrate rapidly compared to viability decay (days), motivating a quasi–steady approximation for *E* and *ε*. (iii) Strains are genetically identical and differ only in physiological state (set by prior growth), resulting in different *β*_0,*i*_, *k*_*i*_, and *γ*_0,*i*_.

### Extracellular energy balance

We denote by *E*(*t*) the amount of energy available in the supernatant, shared by all cells. Energy is supplied when cells die and release recyclable nutrients, and it is consumed when viable cells take up these nutrients.

If *N*_*i*_(*t*) is the number of viable cells of strain *i*, then *dN*_*i*_*/dt <* 0 during starvation. We write the death flux as 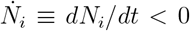. Each dying cell releases a usable amount of energy *α*_*i*_ (the recycling yield). Since 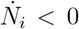, the release term contributing to *dE/dt* enters as 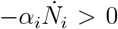. Under the pool-limited (unsaturated) uptake assumption, uptake is proportional to both the number of viable cells and the external pool *E*, with rate constants *k*_*i*_. The extracellular balance is therefore

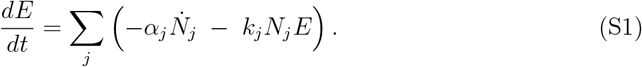

### Intracellular energy balance

Each viable cell maintains an internal energy pool *ε*_*i*_(*t*) (accessible energy per cell). Uptake is proportional to the external pool, with rate *k*_*i*_*E*, while consumption depletes internal energy at a rate *β*_*i*_(*ε*_*i*_), the consumption rate as a function of internal energy:

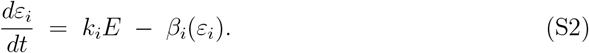

### Quasi–steady state assumption

External and internal energy pools usually equilibrate rapidly compared to viability decay (days), so we set *dE/dt* = 0 and *dε*_*i*_*/dt* = 0. Setting Eq. S1 to zero gives

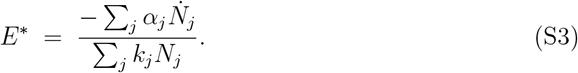

From Eq. S2, the intracellular balance at steady state reads

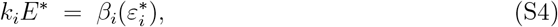

which is the relation quoted in the main text as Eq. (3). Substituting Eq. S3 into Eq. S4 yields

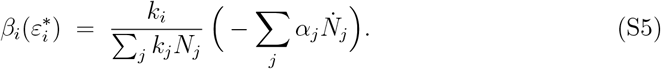

Thus, the consumption rate of strain *i* equals its share of the uptake capacity, 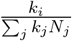 multiplied by the total nutrient release, 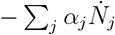.

### Consumption and internal energy

Let 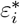 be the quasi–steady internal energy of a cell of strain *i*. A cell with no internal energy cannot consume, while with sufficient energy it can consume what it takes up. We therefore write

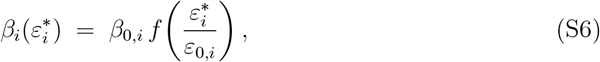

where *β*_0,*i*_ is the reference consumption rate in monoculture, *ε*_0,*i*_ the reference internal energy, and *f* a dimensionless increasing function with *f* (0) = 0 and *f* (1) = 1. Linearizing for small arguments gives

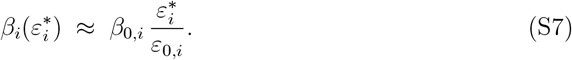

Equating Eq. S7 with Eq. S5 allows us to solve for the internal energy,

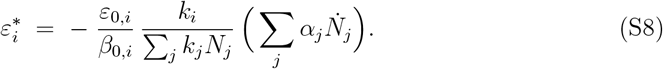

### Death rate and internal energy

We assume that the death rate depends inversely on internal energy:

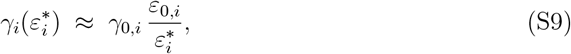

where *γ*_0,*i*_ is the strain–specific reference death rate in monoculture. This is the functional dependence referenced in the main text (Eq. (7)). Substituting Eq. S8 into Eq. S9 gives

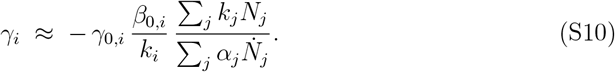

Using 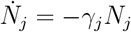 makes the denominator positive, so *γ*_*i*_ *>* 0.

### Coupled equations for death rates

Population decline is linked to death rate by 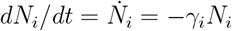(negative), with *γ*_*i*_ *>* 0 by definition. Substituting 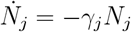 into Eq. S10 yields the coupled implicit relation

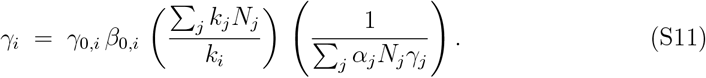

Rearranging Eq. S11 produces, for general multi–strain communities, a quadratic equation in *γ*_*i*_:

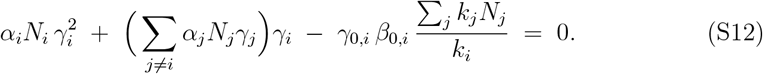

The quadratic formula then gives

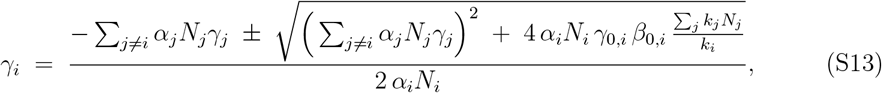

where the “+” branch ensures *γ*_*i*_ *>* 0.

### Compact explicit form for the coupled death rates

From the implicit coupled relation (SI Eq. S11), we introduce

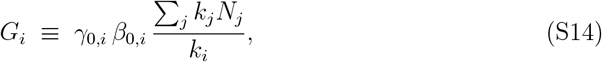

and the community balance

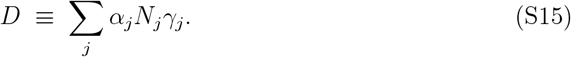

With these definitions the death rate of strain *i* takes the explicit form

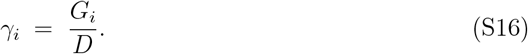

Self–consistency follows by substituting (S16) into (S15), which gives

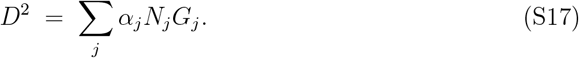

Using the monoculture identity *γ*_0,*i*_ = *β*_0,*i*_*/α*_*i*_, the term *α*_*j*_*N*_*j*_*G*_*j*_ simplifies to 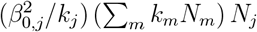. Hence, for implementation it is convenient to write

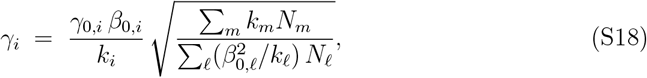

which no longer contains *α* explicitly. This community prefactor is the same one that appears in the two–strain expressions of the main text (Eqs. (9)–(10)).

#### Numerical procedure used for the main text (alpha-free)

At each time increment Δ*t*:

1. Compute *K* ≡∑_*j*_ *k*_*j*_*N*_*j*_.
2. Compute 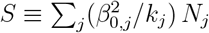.
3. For each strain *i*, evaluate

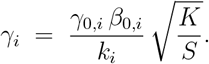
4. Update viabilities:

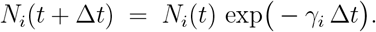

This scheme uses only {*γ*_0,*i*_, *β*_0,*i*_, *k*_*i*_}, as in the main text, and is algebraically equivalent to the coupled system underlying SI Eq. S11.

### Dependence on population ratio for two strains

To gain further intuition, consider two competing strains with fixed physiological parameters (*β*_0,*i*_, *γ*_0,*i*_, *α*_*i*_, *k*_*i*_). Let *N*_1_ and *N*_2_ denote their viabilities and define the viability ratio

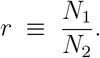

From Eq. S16 together with Eq. S17, each death rate can be written as

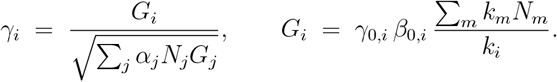

Separating the community-dependent factors, define

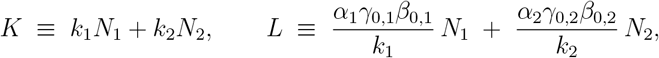

so that

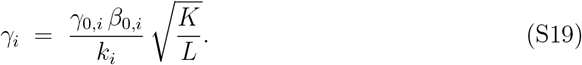

Inserting *N*_1_ = *rN*_2_ and canceling *N*_2_ yields

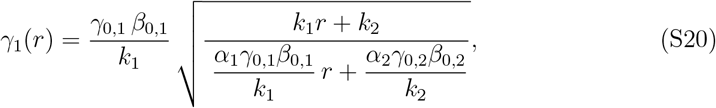

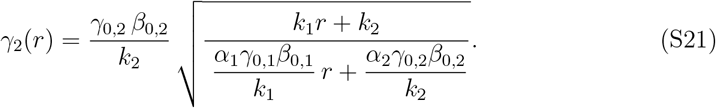

Using *γ*_0,*i*_ = *β*_0,*i*_*/α*_*i*_ eliminates *α*_*i*_ from the denominators, giving the two–strain forms cited in the main text (Eqs. (9)–(10)):

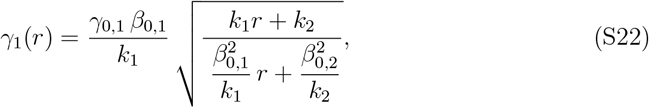

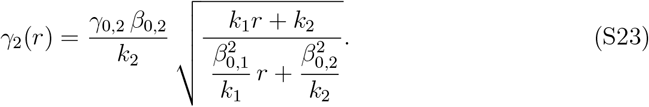

#### Properties

- Both death rates share the same community-dependent prefactor 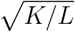, which varies with *r*.
- Their ratio is independent of *r*:

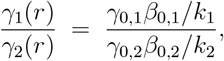

set purely by physiological parameters.
- In the monoculture limits (majority limits), the model recovers reference behaviors: *r* → *∞* ⇒ *γ*_1_ → *γ*_0,1_, *r* → 0 ⇒ *γ*_2_ → *γ*_0,2_.
- In the minority limits, the rare strain’s death rate approaches a constant determined by the physiology of the majority. For strain 1 in the extreme minority (*r* → 0), using Eq. (S22),

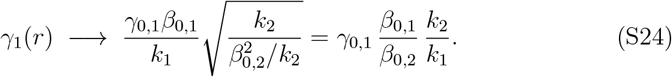

Analogously, for strain 2 in the extreme minority (*r* → ∞),

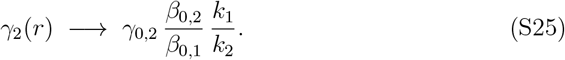

Thus, the absolute magnitude of *γ*_*i*_ depends on the population ratio through the shared community prefactor, whereas the competitive advantage between strains is fully determined by intrinsic physiological parameters.

### Experimental scalings of model parameters

We now connect the model to experimental measurements by inserting growth–rate–dependent scalings for key physiological parameters reported by Biselli et al. [4] for *E. coli* as a function of prior growth rate *µ* (units h^*−*1^). Specifically, we use the dependencies of the consumption rate *β*_0_ and the cell surface area *S* on *µ*, and assume *k* ∝ *S*. These empirical dependencies are well described by exponentials in *µ*, which enables analytical predictions for how death rates change with growth history.

**Exponential scalings from Biselli et al. [4]** Based on the reported fits, we write

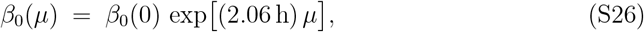

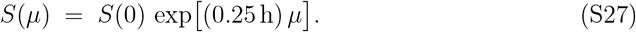

For the surface scaling, we followed Biselli et al. by calculating surface area from cell dimensions (cylinders with hemispherical caps). Implementing *k* ∝ *S* gives

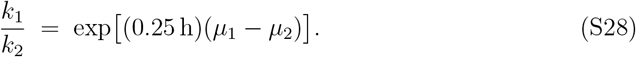

**Analytical results in the limits**. With Eqs. (S26) and (S28), the minority-limit relation (Eq. (14)) becomes the exponential scaling quoted in the main text (Eq. (15)):

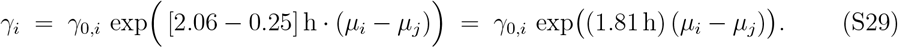

For the slowest (mannose) and fastest (glucose) growth conditions, *µ*_man_ ≈ 0.22 h^*−*1^ and *µ*_glu_ ≈ 0.90 h^*−*1^, this yields

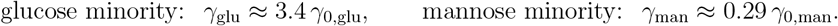

These limiting predictions provide a compact, intuitive rule: minority death rates scale exponentially with the difference in prior growth rates, reflecting the combined effects of maintenance demand and uptake share encoded in Eq. (14).

## B Supplementary Tables

**Table S1:** Growth and death rates of strains labeled with chloramphenicol (Cm) or kanamycin (Km) resistance markers, grown on minimal medium with either mannose, glycerol, or glucose. No significant differences were observed for any growth substrate (paired t-test, two-tailed).

| Metric | Substrate | Cm | Km | p-value | n |
| --- | --- | --- | --- | --- | --- |
| Growth rate<br>(h<sup>-1</sup>) | mannose | 0.21 ± 0.01 | 0.22 ± 0.02 | 0.24 | 3 |
|  | glycerol | 0.715 ± 0.002 | 0.72 ± 0.01 | 0.59 | 2 |
|  | glucose | 0.88 ± 0.04 | 0.92 ± 0.01 | 0.25 | 3 |
| Death rate<br>(h<sup>-1</sup>) | mannose | 0.0149 ± 0.0001 | 0.015 ± 0.002 | 0.69 | 3 |
|  | glycerol | 0.020 ± 0.003 | 0.0200 ± 0.0003 | 0.89 | 2 |
|  | glucose | 0.025 ± 0.002 | 0.025 ± 0.001 | 0.59 | 3 |

## Notes

### Competing Interest Statement

The authors have declared no competing interest.

### Summary of Updates

We corrected an error in the author list.

